# Directional selection failed to produce changes in olfactory Y-maze learning and memory but induced changes in climbing phenotypes

**DOI:** 10.1101/2025.05.18.654242

**Authors:** Victoria L. Hamlin, Jamie Baumann, Reiley Heffern, Alaina Franke, Elizabeth G. King

## Abstract

Learning and memory are fundamental complex traits that allow for assessment and response to changes in their environment. Beyond cognition, these high order traits require several subcomponents, from sensory perception to motor output, in order to execute the intended response to a stimulus. Within the population, we see variation among individuals in abilities to perform learning and memory tasks. It is still largely unknown what genetic factors contribute to variability in these phenotypes; therefore, our study aims to gain better insight by utilizing a directional selection paradigm to drive differences in olfactory learning and memory behavior. Directional selection experiments allow for evaluation of the response to selective pressures across multiple biological levels through amplification of phenotype differences between groups. We used a reward based olfactory associative learning and memory assay to train a synthetic population of flies allowing only those who passed both tests to mate across ten generations. Our study shows significant changes in the climbing subcomponent required to perform well on the y-maze assay, however, we did not observe any significant changes to olfactory learning and memory behavior.

## 1 Introduction

Directional selection has often been used to amplify phenotypic differences between control and experimental lines with the aim of measuring the response to selective pressure across multiple biological levels (Harshman & Hoffmann 2000; Garland & Rose 2009). At the genomic level, allele frequencies can be compared between groups to identify correlations with changes seen at the phenotypic level, providing potential insight to critical gene networks involved in direct and indirect traits affected by the applied selective pressure (Burke & Rose 2009; Garland & Rose 2009). When applying stressors to a genetically diverse population these changes can happen rapidly in sexually reproducing organisms (Gibbs 1999; Burke *et al*. 2010) in which some studies show changes in as little as ten generations (Hoffmann & Parsons 1993; Swallow *et al*. 1998; Mery & Kawecki 2002).

*Drosophila melanogaster*, fruit flies, are well suited for selection and experimental evolution work due to the ease of maintenance, short timespan between generations, wealth of community tools, and ability to generate large population sizes (Burke & Rose 2009; Burke *et al*. 2010). Directional selection projects have been done with *D. melanogaster* on a range of phenotypes including starvation resistance (Chippindale *et al*. 1996), aging (Stearns 2000), and post-infection survival rates (Basu *et al*. 2024). Behavioral traits have also been shown as modifiable through directional selection experiments. Increases in learning speed and longer term memory were induced using an assay where oviposition substrate preferences were modified by pairing with a bitter substance (Mery & Kawecki 2002). Selection for different levels of aggressive behaviors in male flies was able to produce fly lines exhibiting higher or lower aggression in comparison to controls (Edwards *et al*. 2006). Recent studies have provided evidence that even extended phenotypes, such as social network positions which require measurements of interaction with conspecifics, have low-to-mid range broad sense heritabilities and are likely to undergo directional selection (Wice & Saltz 2021).

Indirect and unanticipated mechanisms and traits have also been shown to be affected by selection experiments highlighting the importance of measuring many aspects and potential sub-characteristics of complex multifactorial traits (Harshman & Hoffmann 2000). Indirect selection on pre-adult development time was induced by experimental design when females in one experimental group had a selective pressure to lay fertile eggs at 14 days post oviposition while the second group had 8 additional weeks before egg collections were conducted (Chippindale *et al*. 1994). In an experiment aiming to increase ethanol tolerance, a subset of selected lines also developed increased tolerance to acetone, a molecule not directly related to ethanol metabolism (Hoffmann & Cohan 1987) indicating that multiple mechanisms are being acted upon in response to the selective pressures placed on the population. Overall, these prior studies using directional selection have provided the field with insight on the dynamic responses to environmental pressures from the genomic to behavioral level. In addition to improving the understanding of trait evolution, these experiments have provided insight to the complex trade-offs between phenotypes, co-evolution of correlated traits, and the diversity of adaptive measures that can evolve under the same pressure (Harshman & Hoffmann 2000; Swallow & Garland 2005; Garland & Rose 2009).

Olfactory learning and memory are complex high order traits comprised of many sub-phenotypes that are influenced by multiple genetic loci, environmental factors, and complex interactions between the two. As characteristic of a complex trait, there is a spectrum of abilities in olfactory learning and memory tasks between different genetic backgrounds as identified in a previous study conducted in our lab (Hamlin *et al*. 2025). Utilizing a y-maze olfactory appetitive learning and memory assay, we also identified variability in subcomponents of this assay including odor preference, odor acuity, sugar acuity, and climbing abilities (Hamlin *et al*. 2025). The genetic mechanisms underlying the phenotypic variability displayed in these phenotypes are still vastly unknown. With the ability to manipulate both direct and indirect mechanisms, selection experiments have proven to be a valuable tool to uncover the dynamic interplay of the many mechanisms at work to produce complex phenotypes. We hypothesize that use of a directional selection paradigm to impose change in olfactory learning and memory behavior can provide further insight into the genetic variants influencing the phenotypic outcomes. To test this, we perform a directional selection experiment utilizing an olfactory associative learning and memory Y-maze assay on a synthetic population of flies with the aim to increase performance outcomes in the experimental lines. Additionally, we analyze the sub-characteristics contributing to the overall learning and memory phenotype which include odor preferences, climbing, and performance in different light sources.

## 2 Methods

### Creating a Synthetic Population

The genomic variation in the starting population is important for selective pressures to influence changes in allele frequencies. We created two synthetic populations by placing ten mated females from 97 recombinant inbred lines (RILs) from the *Drosophila* Synthetic Population Resource (DSPR) into each cage (Table 1). One week following the cage creation, an egg collection plate was added, and females were allowed to lay eggs for 24 hours. Eggs were then collected by removing a small thin layer of the food plate and placed into a fresh food vial. This step was repeated allowing for two collections per cage. At 13 days post oviposition, the flies from each collection were transferred to a new cage – now 4 cages total – and allowed to randomly mate for 6 days. On day 6, a fresh egg collection plate was added to each cage and eggs collected 24 hours later; the offspring (Generation 2) of this generation (Generation 1) will be the first to have a mix of RIL genotypes. From Generation 2 onward, we began to add yeast paste in addition to food plates in the cages to increase egg counts allowing for multiple egg collections from each cage. From the Generation 3 cages, we doubled the collection from each cage to start the four experimental lines which undergo selection for 10 generations.

**Table 1.**
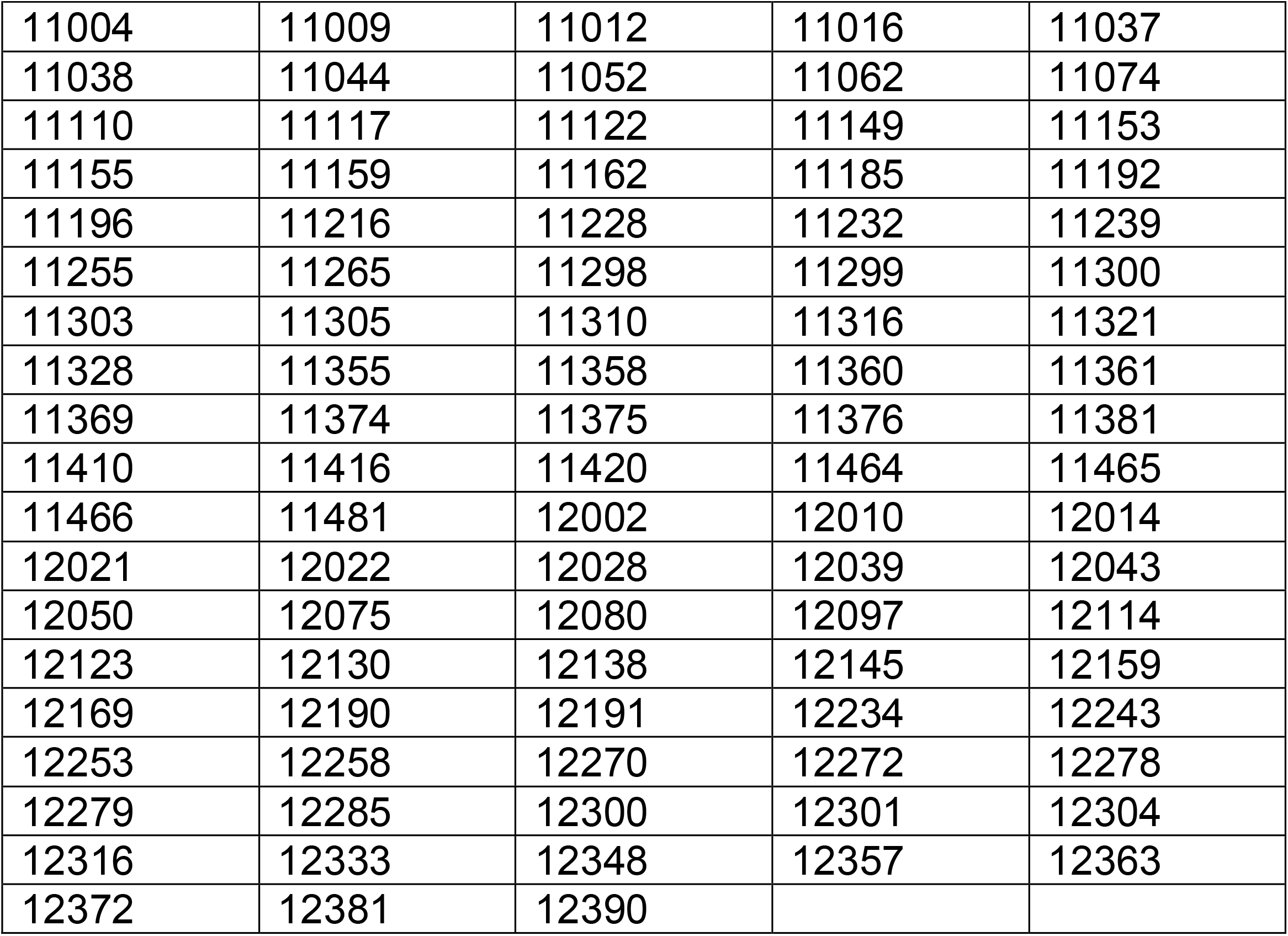
List of DSPR RILs used to create synthetic population. 10 mated females from each RIL were used to start the population.

### Conditioning Assay

To select on olfactory learning and memory phenotypes we use an appetitive conditioning behavioral assay in which flies are trained to associate specific odors with a sucrose reward in mixed sex groups. Initially, flies are starved for 18-20 hours prior to the start of conditioning by transferring to a vial with non-nutritional agar to increase motivation to seek food. The sucrose and non-nutritional agar vials used for the conditioning assay were placed in fly cages the night prior to conditioning to “pre-fly” the vials so the scent of other flies is present during conditioning. Odor caps containing 20uL of either 3-Octanol (OCT) or 4-Methylcyclohexanol (MCH) were placed on the vials 20 minutes prior to the start of the assay to allow for odorization. The conditioning protocol has four sequential phases: (1) flies are transferred into the negative conditioned stimulus (CS-) vial containing non-nutritional agar for five minutes, (2) flies are transferred into an odor free empty vial for five minutes, (3) flies are transferred to the positive conditioned stimulus (CS+) vial containing 2M sucrose agar for five minutes, and then (4) flies are transferred to an empty vial and allowed at minimum five minutes of rest before beginning the testing phase. Half of the vials were trained to associate sucrose with OCT and the remaining half trained to MCH. In our study we tested all generations on a standardized concentration of 0.06% OCT and 0.09% MCH diluted in mineral oil as done in a previous study conducted in our lab (Hamlin *et al*. 2025).

### Y-maze Assays

We conducted learning and memory testing in 3D printed Y-mazes (Mohandasan *et al*. 2022), which are made of 3 chambers; two odor chambers in the top arms and a starting chamber at the base. 1000µl pipette tips were trimmed at the narrow end to make an opening large enough for the flies to enter the chamber but did not allow them to exit, making their initial choice permanent. Y-mazes are odorized for 20 minutes prior to the start of the assay by placing filter paper with 20µL of diluted odorant (odor card) into its respective odor chamber. Flies from the final vial of the conditioning assay were transferred into the starting chamber then placed in a slot under a fabric lightbox chamber and allowed 30 minutes to select between the two odors presented during conditioning. All flies were then collected and categorized based on their choice of either CS+, CS-, or no choice (NC) for those who remained in the starting chamber at the end of the assay. Flies who selected the sucrose paired odor (CS+) were tested for memory recall 4 hours after completion of the conditioning assay using the same Y-maze set up. Flies were then collected based on the choices of CS+, CS-, and NC and those who selected the sucrose paired CS+ odor in both the learning and memory Y-maze assays were released into a cage. Flies that were trained to OCT were released into the same cage as those trained to MCH to try to eliminate any selection on odor preference. Each cage contained Bloomington’s Nutrifly (“Bloomington Drosophila Stock Center” 2025) food and water in addition to supplemental yeast for the first two days to increase egg production. Yeast was removed and a fresh food plate for egg collection was placed in the cage for approximately 24 hours before removal. A new egg collection plate was placed in the cage and allowed a second 24-hour period for a second egg collection. Eggs were collected by taking small portions of the top layer of food and transferring them into a small food vial. Vials were placed in a temperature control chamber for 16 days after the last egg collection then prepared for conditioning and testing as described above for ten generations. After all egg collections were completed, flies were frozen and stored for later DNA sampling. For the first 7 generations of selection the light source used above the lightbox was white light, however, at generation 8 we began to use a red-light source above the lightbox because we speculated that phototaxis may have been overriding the learning and memory phenotype.

To analyze the learning and memory performance across generations, flies were individually scored based at two choice points using a binomial scoring method (Hamlin *et al*. 2025); the climbing choice point where they are scored on whether they enter an odor chamber and the correct choice where they are scored on whether they selected the sucrose paired odor. We took cumulative scores for each cage by calculating the proportions at the climbing and correct choice points. Additionally, we calculated an overall proportion for the selected and control lines at each generation.

### Odor Preference Measurements

Odor preference measurements were conducted in the Y-mazes with unconditioned flies from generations 7 and 9. Flies were transferred into the starting chambers of Y-mazes containing OCT and MCH in the upper chambers of the maze. Y-mazes were placed under the fabric lightbox chamber and flies were allowed 30 minutes to select between the two odors. All flies were then collected and categorized based on their choice of either OCT, MCH or NC. Flies were individually scored based at two choice points using a binomial scoring method (Hamlin *et al*. 2025); the climbing choice point which is scored the same as above and the odor chamber choice where they are scored on if they selected OCT or not. Since the odor chamber choice is scored in relevance to OCT a high score means strong OCT preference while a low score means stronger MCH preference. We also took cumulative scores for each cage as well as line type by calculating the proportions at the climbing and odor choice points.

### Odor Acuity Measurements

Odor acuity measurements were conducted in the Y-mazes with unconditioned flies from generation 10. Flies were transferred into the starting chambers of Y-mazes containing one of the trained odors, OCT or MCH, in one upper chamber of the maze while the other chamber did not contain an odor card (Null Chamber). Y-mazes were placed under the fabric lightbox chamber and allowed 30 minutes to select. All flies were then collected and categorized based on their choice of either Odor Chamber, Null Chamber or NC. Flies were individually scored based at two choice points using a binomial scoring method(Hamlin *et al*. 2025); the climbing choice point which is scored the same as above and the chamber choice where they are scored on if they selected the odor chamber or null chamber. We also took cumulative acuity scores for each cage as well as line type by calculating the proportions at the climbing and chamber choice points.

### Red Light Baseline Odor Preference Assay for RILs

For this experiment, we chose nine RILs from the starting population that were previously identified as potential high performers in a study conducted in our lab (Hamlin *et al*. 2025). To establish each RIL’s baseline preference for OCT versus MCH, flies were placed directly into the Y-maze without prior odor conditioning and allowed to choose between the two odors for 30 minutes. This process was conducted on all RILs in both red and white light. Flies from the OCT vial, the MCH vial, and the NC vial were collected and counted. A preference score was calculated for each RIL for both light conditions by taking the overall proportion of flies choosing OCT, therefore, scores closer to one indicate an OCT preference and scores closer to zero indicate a MCH preference.

### Red Light Learning and Memory Assay for RILs

Following the same conditioning protocol as the selection experiment, we tested learning and memory performance for nine RILs in red light conditions using independent mixed sex groups of flies for learning and memory. Flies allotted for memory testing were put back onto non-nutritional agar for the four-hour waiting period to maintain access to water, while flies for the learning test were tested immediately after conditioning was complete. Flies were then counted based on the vial they were collected from at the end of the 30-minute period Y-maze period. For each RIL, we calculated proportion scores at two decision points: a *climbing* score which measures the proportion of flies leaving the starting chambers and enters either of the upper arms of the Y-maze and a *correct* score which measures the proportion of flies choosing the sucrose paired odor. These measurements were then compared with scores from performance in white light collected in our previous study (Hamlin *et al*. 2025).

### Isofemale Lines Establishment

After completion of the 10^th^ generations’ learning and memory assay, twenty mated female flies from each replicate cage were used to create isofemale lines; ten females from the OCT trained group and ten females from the MCH trained group in generation 10. Each individual female was placed into a food vial to ensure only her offspring are present. In addition, we collected 20 females of each replicate cage of the control groups and placed females into individual vials for generation of isofemale lines. It is assumed these offspring are half-siblings since female flies can lay eggs from multiple matings at one time. These lines are continually inbred for each generation with efforts to preserve any genomic changes that have occurred over the selection period.

### Isofemale Climbing Assays

Isofemale lines were duplicated and tested for climbing abilities in single vial and Y-maze assays for 5 isofemale lines from each cage and line type, 40 lines total. For the single vial assay, flies were transferred into an empty food vial with a pre-marked threshold line 6.4 cm from the bottom of the vial. After transfer, flies were given at least 2 minutes to rest before undergoing the climbing assay. After the rest period, flies were moved to a white lightbox chamber to record climbing phenotype after the vial has been tapped hard to move all flies to the bottom. After 30 seconds, the recording was stopped, and flies were counted under CO2 knock out to obtain the total number of flies in the assay. Recordings were then counted at the 10 second post tap mark to measure the distribution of flies above the threshold and a proportion score was calculated. This test was also conducted in red light conditions with a new set of flies.

Additionally, we tested climbing abilities in Y-mazes to see if it differs from single vial climbing in red light. Unconditioned flies were transferred into the starting chamber of a Y-maze with no odors. Flies were allowed 20 minutes to move through the Y-maze and were collected based on final chamber location. Flies were categorized into an upper chamber count for those who were collected from either of the two upper chambers, and a lower chamber count for those who did not leave the starting chamber.

### Heritability Calculation for Learning, Memory, and Climbing Phenotypes

Using data collected for 50 DSPR RILs on olfactory learning and memory from previous work in the lab(Hamlin *et al*. 2025), we calculated the broad-sense heritability estimate of olfactory learning, memory, and climbing from both rounds in the Y-maze assays. The heritability estimates measure the proportion of the variation in phenotype due to genetic verses environmental factors with scores closer to 1 being more strongly influenced by genetic factors and scores closer to 0 being more strongly influenced by environmental factors. For each trait, we estimated the genetic and phenotypic variance and covariance components from a linear mixed model using the *lme* and *VarCorr* functions in the *nlme* package (Pinheiro & Bates 2000; Pinheiro J, Bates D, R Core Team (2024) n.d.). Prior to fitting this mixed model, we quantile normalized each trait to ensure normality.

### Data Availability Statement

Raw data is available from zenodo (https://zenodo.org/): 10.5281/zenodo.15425618. Founder genotype assignments from the hidden Markov model are available as two data packages in R (http://FlyRILs.org/) and are available from the Dryad Digital Repository (http://dx.doi.org/10.5061/dryad.r5v40). See King et *al*. (King *et al*. 2012b, 2012a) for details of the DSPR datasets. The complete code used to perform all analyses is available at GitHub (https://github.com/EGKingLab/OlfactorySelection).

## 3 Results

### Heritability of Phenotypes

Heritability was calculated using data collected on 50 recombinant inbred lines that contributed to the base population used in the selection experiment. The calculated heritability estimates based of the correct proportion scores are 0.069 for olfactory learning and 0.103 for olfactory memory. Additionally, we calculated the heritability of the climbing phenotypes in the learning and memory rounds based of the proportion of flies leaving the starting chamber, regardless of which odor chamber was selected. The heritability estimate for climbing in the learning and memory rounds are 0.61 and 0.305 respectively.

### Selection Results for Learning & Memory

Selection on olfactory learning and memory phenotypes were conducted for ten generations on a synthetic population of flies with founders from 97 DSPR RILs (Table 1). Scores for olfactory learning and memory were measured for both selected and control groups at generations 4, 5, 7, 9, and 10. The overall pattern across the testing period showed limited changes in performance of selected groups over time indicating that selection on olfactory learning and memory was ineffective (Figure 2). Comparing the fourth and last generation scores for learning, our selection mean proportion score across all cages is 0.522 at generation 4 and 0.521 at generation 10. For memory, the mean proportion score across all cages is 0.473 for generation 4 and 0.542 for generation 10. While this does increase, the final score at generation 10 is not significantly different from the control group which has a score of 0.587 (Table 2).

**Table 2.** Comparison of mean proportion scores across generations for each phenotype tested. Phenotypes were compared at Generations 4, 5, 7, 9, and 10. Adjusted p-values with asterisk indicates significant difference between controls and selected groups.

| **Generation** | **Selection** | **Control** | **Adjusted P-Value** |
| --- | --- | --- | --- |
| *Learning Proportion Score* |  |  |  |
| 4 | 0.522 | 0.584 | **2.63e<sup>-4</sup> |
| 5 | 0.541 | 0.538 | 1.00 |
| 7 | 0.513 | 0.554 | 0.393 |
| 9 | 0.516 | 0.563 | *0.039 |
| 10 | 0.521 | 0.508 | 1.00 |
| *Memory Proportion Score* |  |  |  |
| 4 | 0.473 | 0.567 | **3.42e<sup>-3</sup> |
| 5 | 0.473 | 0.516 | 0.393 |
| 7 | 0.480 | 0.403 | 0.289 |
| 9 | 0.460 | 0.534 | 0.376 |
| 10 | 0.542 | 0.587 | 0.289 |
| *Climbing – Learning Round* |  |  |  |
| 4 | 0.885 | 0.900 | 0.393 |
| 5 | 0.906 | 0.879 | **5.34e<sup>-3</sup> |
| 7 | 0.896 | 0.798 | ***7.30e<sup>-16</sup> |
| 9 | 0.685 | 0.542 | ***1.47e<sup>-32</sup> |
| 10 | 0.749 | 0.517 | ***5.03e<sup>-47</sup> |
| *Climbing – Memory Round* |  |  |  |
| 4 | 0.825 | 0.857 | 0.289 |
| 5 | 0.874 | 0.884 | 1.00 |
| 7 | 0.882 | 0.803 | **8.59e<sup>-5</sup> |
| 9 | 0.749 | 0.689 | 0.064 |
| 10 | 0.717 | 0.708 | 1.00 |

**Figure 1.**
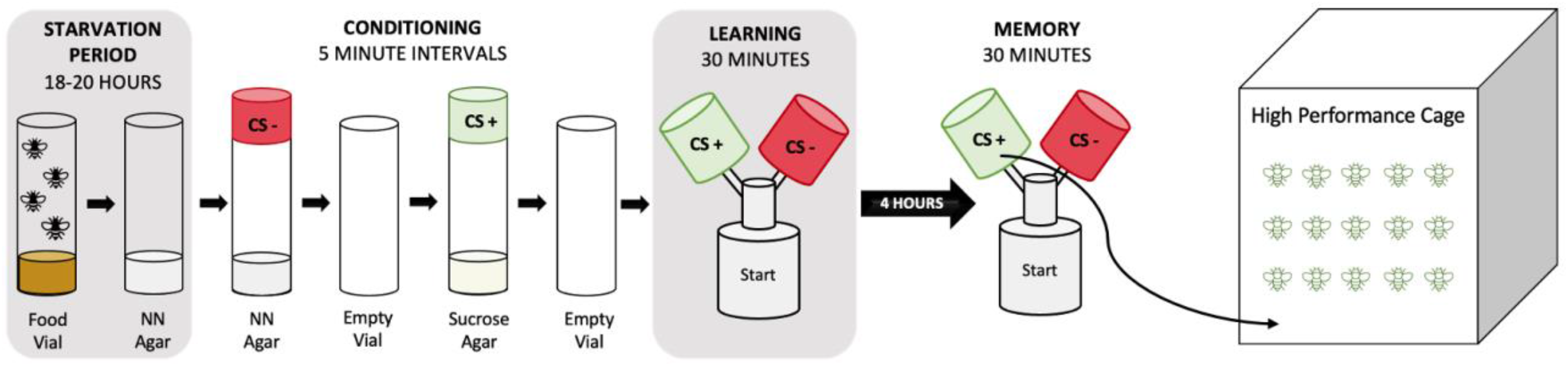
Schematic of Selection Process. Approximately 60-80 flies are placed in a vial with non-nutritional agar (NN Agar) for an 18-to-20-hour starvation period. Flies are conditioned by exposure to the negative conditioned stimulus (CS-) in vials with NN agar then allowed to rest for 5 minutes. Flies are transferred to the positive conditioned stimulus (CS+) vial which contains 2M sucrose agar then allowed to rest again before transfer into the starting chamber of the Y-maze. Flies are given 30 minutes in the y-maze to choose between the two odors used during conditioning. The learning assay is conducted immediately after conditioning then 4 hours later the memory assay is conducted on flies who selected the CS+ odor during the learning phase. Flies who choose the sucrose paired chamber during the memory assay are transferred to a cage where they are allowed to randomly mate.

**Figure 2.**
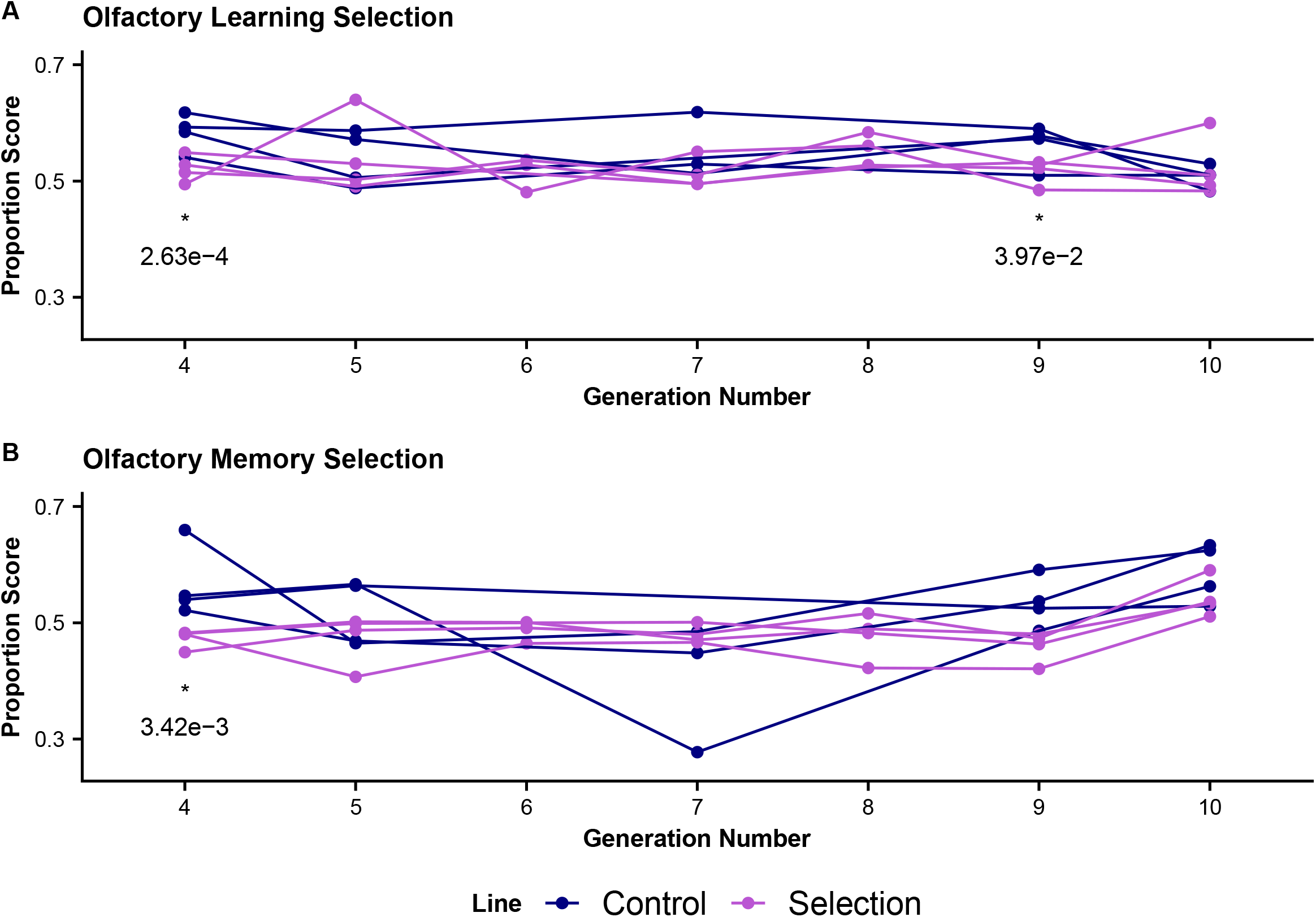
Olfactory Learning and Memory Selection Trends. The proportion of flies choosing the sucrose trained odor is plotted for generations 4 through 10. Testing in generation 1-7 were conducted using a white light source while generation 8-10 are conducted in red light indicated by shading in graph. **(A)** Learning Assay scores for control (blue) and selection (magenta) lines on the learning assay. We see all groups tend to have similar scores across all generations. Marginally significant differences are found at generations 4 and 9 **(B)** Learning Assay scores for control (blue) and selection (magenta) lines on the learning assay. We see all groups tend to have similar scores across all generations. Marginally significant differences are found at generation 4.

We used a generalized linear mixed model to analyze differences in performance between selected and control groups with the individual binomial scored data. We found in the learning assay, generations 4 (adjusted p < 0.001) and 9 (adjusted p = 0.039) show a significant difference between control and selected lines in which controls are performing better than the selected group (Figure 2A, Table 2). In the memory assay, generation 4 (adjusted p = 0.003) also shows that the control group is performing better than the selected (Figure 2B, Table 2). These significant differences do not persist over all generations. Importantly, the magnitude of the difference in performance is very small, indicating that the significant result is likely due to our high power from our large sample size.

### Odor Preference

To measure if the selective pressure changed odor preferences between lines and generations, we measured a subset of flies from each cage at generation 7 and 9. These flies did not undergo any conditioning and were measured for preference between the two odors used in the y-maze. The mean OCT choice proportion score for each generation and line are 0.347 for the generation 7 selection line, 0.376 for the generation 7 control line, 0.349 for the generation 9 selection line, and 0.336 for the generation 9 control line (Figure 3A). These scores indicate there is a slight preference towards the MCH odor, however, using the individual binomial score data for analysis, we see no significant differences in odor preferences between the two generations tested or between the control and selected lines.

**Figure 3.**
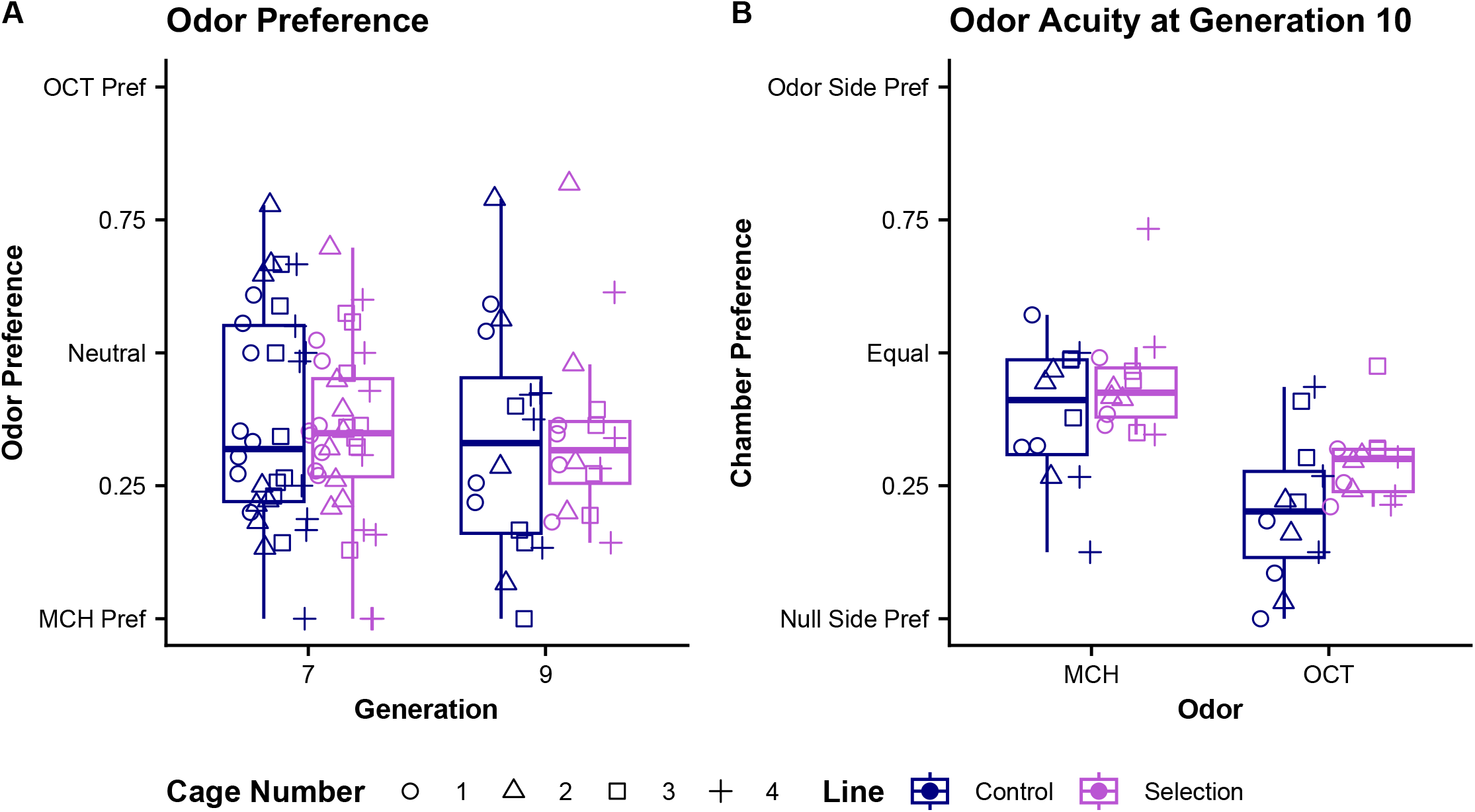
Analysis of odor performance and acuity. **(A)** Odor preference was measured at generations 7 and 9 on untrained flies. There are no significant difference in odor preference between generations or line types **(B)** Acuity measurements were taken at generation 10 for both odors. For MCH there are no significant differences in acuity performance between lines. For OCT there is a marginally significant difference in acuity performance between lines. For comparison between odors, we see a significant difference between MCH and OCT, with OCT having more flies entering the null vs odor chamber.

### Odor Acuity

At generation 10, we measured odor acuity by placing unconditioned flies in Y-mazes which only have one of the two upper chambers containing an odor card. The mean odor chamber choice proportion scores are 0.387 for the control line and 0.446 for the selection line when measured for MCH acuity. When measured for OCT acuity the mean odor chamber choice proportion scores are 0.204 for the control line, and 0.291 for the selected line (Figure 3B). Using the individual binomial scores, we do not see a significant difference between odor chamber choice in the control and selection lines when presented with the MCH odor. However, we see a marginally significant difference in odor chamber choice between control and selected lines when presented with the OCT odor (p = 0.036). We also see a significant difference between odor chamber choice between the two odors regardless of line, with OCT being significantly lower than MCH (p < 0.001).

### Light Source Comparisons of Recombinant Inbred Lines

We measured the effect of light source on performance in the learning, memory, and odor preference assays for a subset of the RILs used to make the base population for selection experiments (Figure 4). Using binomial scored data, we use a generalized linear mixed model to determine if there are significant differences between the performance in red light compared to white light. In the learning assay, light source has a marginally significant impact on performance (adjusted p = 0.005) in which flies perform better in white light across all RILs (Figure 4A). In the memory assay, light source had no significant impact on performance across all RILs (Figure 4B). When measuring odor preference between OCT and MCH, there is no significant difference in performance between light sources (Figure 4C).

**Figure 4.**
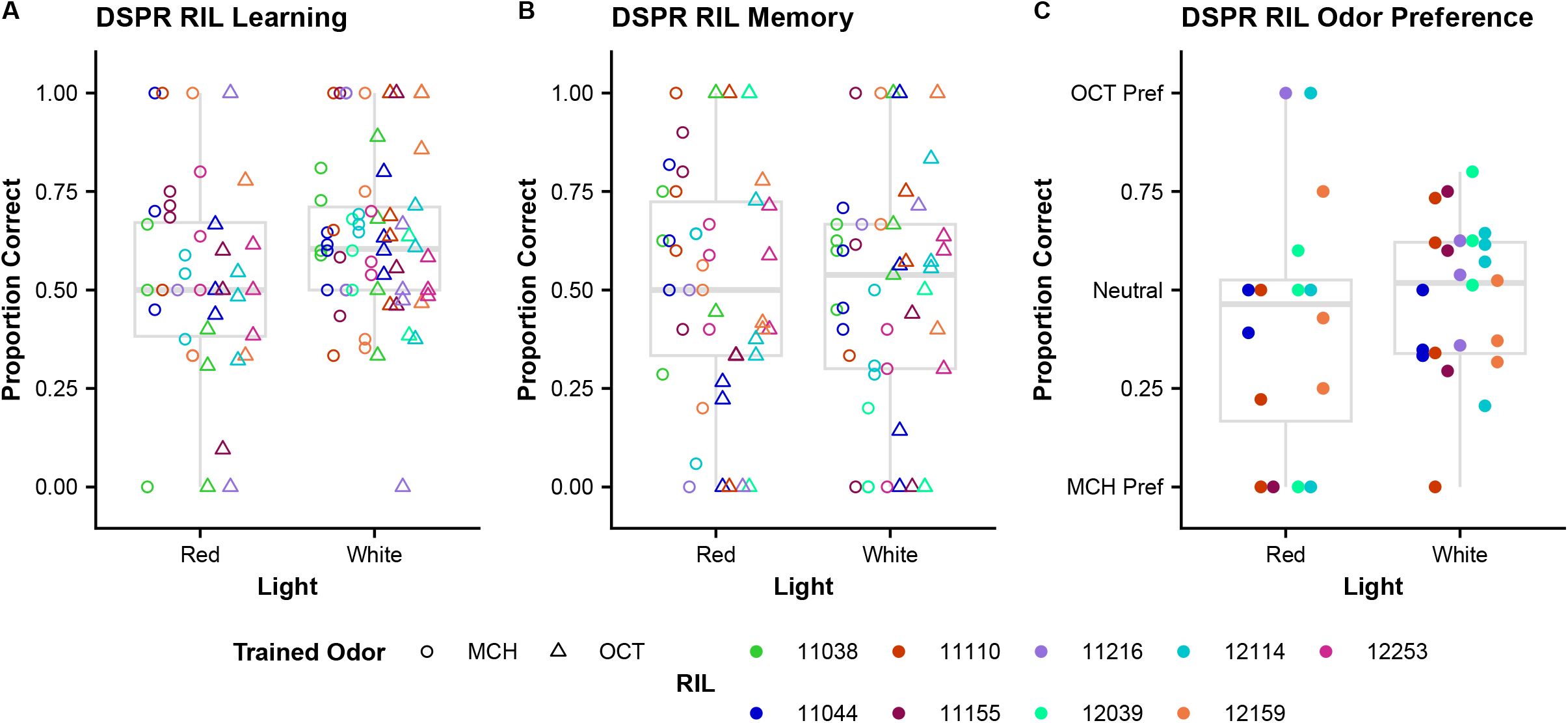
Comparison of DSPR RIL performance in red and white light. **(A)** Difference in performance between RILs in the learning assays conducted in red and white light. Impact of light source is marginally significant (adjusted p = 0.005). **(B)** Difference in performance between RILs in the memory assays conducted in red and white light show no significant difference in performance. **(C)** Odor preference scores across RILs in red and white light conditions are not significantly different.

### Selection on Climbing Phenotype

Using the binomial scoring method, we analyzed the impact of our selection protocol on climbing abilities by scoring flies on if it left the starting chamber or not. Scores for climbing were measured for both selected and control groups at generations 4, 5, 7, 9, and 10 in both learning and memory assays. The overall pattern across the testing period showed significant changes in performance of the selected groups in the learning assay overtime indicating there was selection on the climbing phenotype (Figure 5A). However, we see little difference between the climbing phenotype between control and selected lines during the memory assay (Figure 5B).

**Figure 5.**
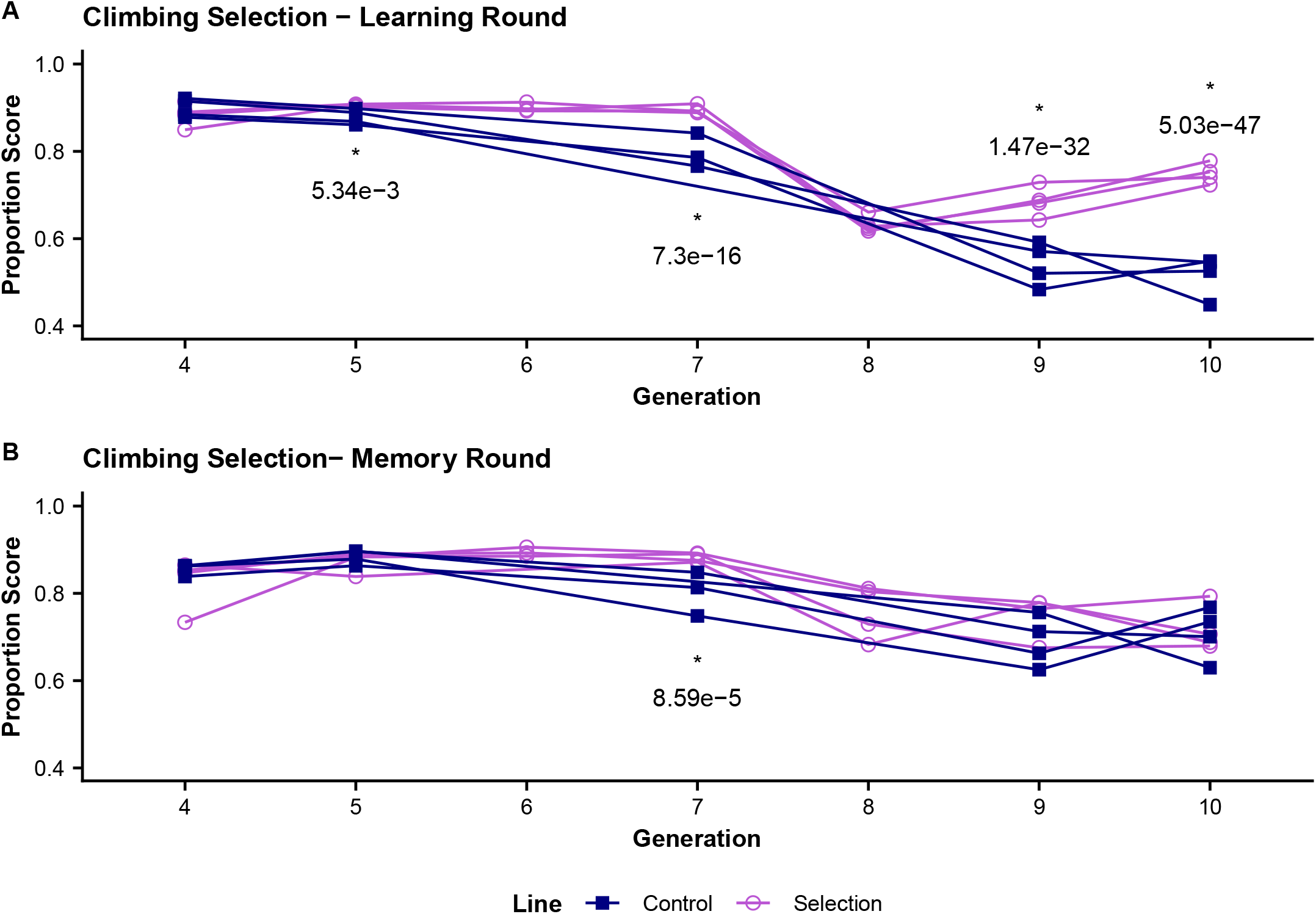
Climbing Control and Selection Trends. The proportion of flies climbing into either odor chamber was measured for generation 4 -10. Testing in generation 1-7 were conducted using a white light source while generation 8-10 are conducted in red light indicated by shading in graph. **(A)** Climbing proportions during the learning assay shows significant changes in the climbing proportion at generations 7, 9, and 10 and a marginal significance at generation 5. **(B)** Climbing proportions during the memory assay are similar across generations with significant difference in performance at generation 7.

Comparing the scores in the first and last measured generations for the learning round, the mean climbing proportion score in generation 4 (white light testing) is 0.885 for the selected lines and 0.900 for controls and in generation 10 (red light testing) 0.749 for selected and 0.517 for controls (Table 2). Comparing the scores in the first and last measured generations for the memory round, the mean climbing proportion score in generation 4 (white light testing) is 0.825 for the selected lines and 0.857 for controls and in generation 10 (red light testing) 0.717 for selected and 0.708 for controls (Table 2).

We again used a generalized linear mixed model to analyze differences in climbing performance between selected and control groups with the individual binomial scored data. The climbing phenotype in the learning round showed significant differences between the control and selected lines at generations 5 (adjusted p = 0.005), 7 (adjusted p < 0.001), 9 (adjusted p < 0.001), and 10 (adjusted p < 0.001) with the selected line outperforming the controls when measuring the proportion of flies entering one of the upper chambers (Figure 5A, Table 2). The climbing phenotype in the memory round only showed significant difference between the selected and control lines at generation 7 (adjusted p < 0.001) while scores for the other measured generations are relatively similar (Figure 5B, Table 2).

### Isofemale Climbing Assays

In the single vial assay, we see a significant difference in climbing proportions between tests conducted in red light in comparison to white light conditions (Figure 6A). The mean climbing proportion scores for each condition and line are 0.271 for red light controls, 0.385 for red light selection, 0.742 for white light controls, and 0.667 for white light selection. Both the selected and control lines have significantly higher climbing proportions during the white light conditions compared to their counterparts tested in red (adjusted p < 0.001). There is no significant difference in climbing proportions between selected and control groups in white light, however, we see a marginally significant difference between selected and control lines in red light with the selected lines having a larger climbing proportion (adjusted p = 0.055).

**Figure 6.**
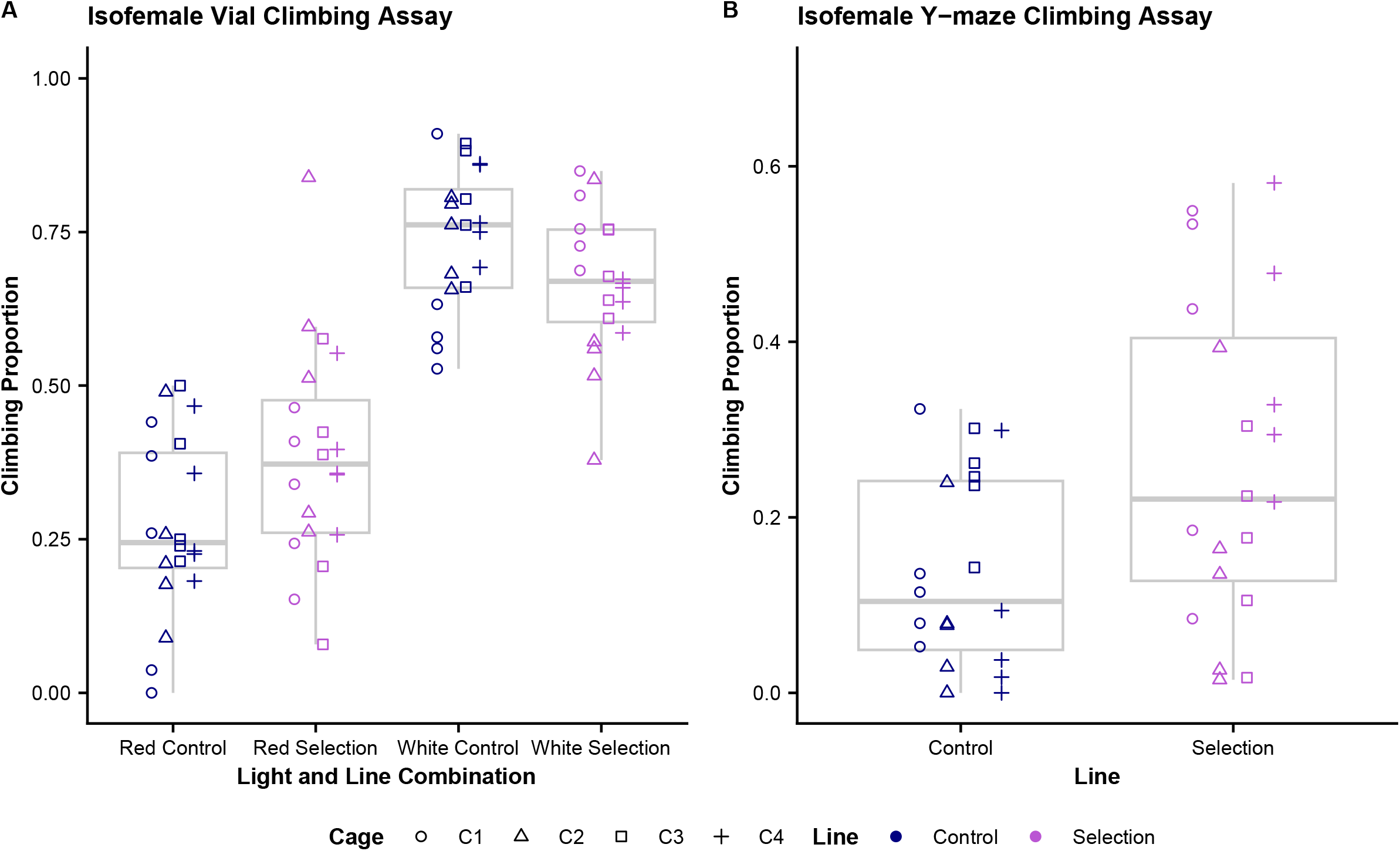
Measurement of climbing performance in Isofemale Lines derived from control and selection cages show the climbing phenotype may be broadly applicable. **(A)** Single vial climbing comparisons between control and selection derived isofemale lines in red and white light. We see there is a significant difference in climbing performance between the two light conditions (adjusted p < 0.001). In red light there is marginal significance in the difference between controls and selected lines, with selected performing slightly better (adjusted p = 0.055). There is no significant difference in white light. **(B)** Isofemale selection and control derived lines show significant difference in performance in the y-maze climbing test, with selection outperforming controls (p <0.001).

Using a generalized linear mixed model, we analyzed the difference in y-maze climbing performance between the control and selected groups with the individual binomial score data. In the y-maze climbing assay, we see a significant difference (p < 0.001) in climbing proportion between the two lines with the selection line having a proportion score of 0.262 and control lines a proportion score of 0.138 (Figure 6B).

## 4 Discussion

Through our selection protocol we were unable to produce meaningful change in olfactory learning or memory phenotypes between the control and selection groups after ten generations of selection (Figure 2). While a previous study aimed at selecting olfactory learning and memory has shown significant differences in under ten generations when using oviposition choice as the measurable outcome (Mery & Kawecki 2002), the short time period used in our study may be too early to detect measurable changes when trying to isolate a trait with low heritability if the selective pressure is not strong. Our heritability estimates for olfactory learning and memory based on data from 50 DSPR RILs contributing to the base population of the selection line was 0.069 and 0.103, respectively, suggesting variation in these traits are largely due to environmental influence, and the response to selection may be slow.

Using the Y-maze assay for olfactory learning and memory testing, it is difficult to fully isolate the learning and memory phenotypes from other factors contributing to performance such as innate odor preference, phototaxis, climbing aptitude, and motivation. A previous study using the Drosophila Genetic Reference Panel (DGRP), shows avoidance/attraction responses to olfactory cues have a low to moderate broad-sense heritability of 0.14 to 0.33 and significant variation in response to different odors (Arya *et al*. 2015). Variation in odor response was also seen in our work wherein there tends to be stronger avoidance of OCT in comparison to MCH in the selection and control lines on the odor acuity test (Figure 3B). We also see variation in odor preferences between OCT and MCH across replicates (Figure 3A) as well as in the subset of RILs tested (Figure 4C). We suspect there is meaningful impact of phototaxis highlighted by the significantly larger proportions of flies climbing to the upper chambers and above thresholds in white light conditions as opposed to red light in all assays we conducted. This result is unsurprising since extensive studies in Drosophila show they exhibit reduced sensitivities to longer wavelength light (de Salomon & Spatz 1983; Yamaguchi *et al*. 2010; Little *et al*. 2019).

While we did not see changes in either of our targeted phenotypes, we were able to see significant changes in the climbing phenotype, specifically within the learning round (Figure 5A). We suspect this is specific to the first round in the Y-mazes due to our selection protocol in which we only carried over flies who selected the correct odor during the learning phase into the memory phase. This makes it likely they were preselected for good climbing before the memory trials. Additionally, we see this difference in climbing phenotype persist into the isofemale lines derived from the selection lines as far as 36 generations after the selective pressure has been removed. This phenotype appears to be applicable across different implementations of climbing as we see a significant, though small, difference in the vial climbing assay specific to red light in addition to it being significant in the Y-maze climbing assay.

Learning and memory are high order phenotypes in which there are many different subcomponents involved in trait presentation ranging from a large network of genes to other behavioral sub-characteristics. Implementation of our selection protocol was able to act upon the climbing phenotype highlighting its importance in performance on this assay, however, the lack of change in our targeted phenotype indicates that a stronger selective pressure may be required in future works aiming to target olfactory learning and/or memory specifically. Since complete isolation of the learning and memory phenotype would be difficult to accomplish due to the many components involved in trait presentation, this stronger selective pressure could potentially be accomplished by multiple rounds of conditioning prior to enter the testing phase.

## 5 Conclusion

Overall, our study highlights the complexity of phenotypic presentation and heritability of olfactory based learning and memory. We show that our selection protocol was successfully able to impose change on climbing phenotype, a behavioral sub-characteristic of the overall targeted trait. This work furthers our understanding in heritability of olfactory learning and memory showing that there is high environmental impact of this high order phenotype with influences from light conditions, odor preferences, climbing abilities. With established iso-female lines and samples stored from each generation of selection, there is room to learn more from this study with future experiments. Sequencing for allele frequency comparison between control and selected lines will allow for identification of potential genomic contributors to the climbing phenotype. RNA sequencing of the isofemale lines, shown to have maintained the increased climbing phenotype, can also be utilized to pinpoint the role of expression changes. Our lab has additional behavior assays that may be of interest to explore with the iso-female lines generated through this work as the phenotypes may have some correlation: [1] an exploration assay which utilizes a two-chamber maze with no food or odor that can indicate if the climbing phenotype is related to exploration tendencies (Elkins 2023) and [2] a flight chamber in which flies are tested for flying abilities against fan-generated wind that can indicate if the climbing phenotype is related to changes in muscle composition (unpublished work from the King Lab).

## Acknowledgements

This material is based upon work supported by the National Institutes of Health under Grant No. R35GM149238.

